# Alopecia areata patients show deficiency of FOXP3+CD39+ T regulatory cells and clonotypic restriction of Treg TCRβ-chain, which highlights the immunopathological aspect of the disease

**DOI:** 10.1101/503961

**Authors:** Fatma N Hamed, Annika Åstrand, Marta Bertolini, Alfredo Rossi, Afsaneh Maleki-Dizaji, Andrew G Messenger, Andrew JG. McDonagh, Rachid Tazi-Ahnini

## Abstract

Alopecia areata (AA) is a hair loss disorder resulting from an autoimmune reaction against hair follicles. T-helper 1 cells are a major contributor to this disorder, but little is known about the role of T-regulatory cells (Tregs) in AA. Here, we analysed the distribution of circulating Treg subsets in twenty AA patients with active hair loss and fifteen healthy subjects by flow cytometry. The Treg suppressor HLA-DR+ subpopulation was significantly reduced in the patients (P<0.001) and there were significantly fewer cells expressing CD39 among the CD4+CD25+Foxp3+ Treg subpopulation in patients (P=0.001). FOXP3 CD39 Treg cells were also reduced in hair follicles; by 75% in non-lesional skin and 90% in lesional skin, when compared to control healthy skin. To further characterise Treg cells in AA; Tregs (CD4+CD25+FOXP3+) were investigated for their TCRβ sequence. PCR products analysed by Next Generation Sequencing techniques, showed that all frequent public clonotypes in AA Tregs were also present in controls at relatively similar frequencies, excepting two public clonotypes: CATSRDEGGLDEKLFF (V15 D1 J1-4) and CASRDGTGPSNYGYTF (V2 D1 J1-2), which were exclusively present in controls. This suggests that these Treg clonotypes may have a protective effect and that they may be an exciting subject for future therapeutic applications.

## Introduction

Alopecia areata is a common disorder of hair and nails with a lifetime disease risk of 2% in the general population. Autoimmunity is considered central to the pathogenesis [1,2]. Multiple major genetic factors have been demonstrated to be aetiologically important, and HLA genes of the major histocompatibility complex are strongly linked with AA, in common with a number of other autoimmune diseases, [1,3,4]. In addition to clinical associations with autoimmune thyroid diseases and vitiligo, there is a particularly high rate of AA in the rare recessive syndrome of type 1 autoimmune polyendocrinopathy (APS1), caused by mutation of the autoimmune regulator (AIRE) gene [5,6]. T-lymphocyte infiltration of hair follicles is a cardinal feature of the histopathology of AA and in recent years, there has been a realisation that Th17 subsets are likely to be significant agents in the disease process. A single previous study reported the possible involvement of Th17 cells in the pathogenesis of AA [7]. It has also been shown that in peripheral blood mononuclear cells (PBMC), Th17 levels were higher in patients with active disease of short duration, whereas Treg levels were lower in those patients with mild AA [8]. However, the authors did not analyse Treg cell subsets in AA. The characterisation of Treg populations in other autoimmune diseases has revealed defects in the Treg suppressor cell subpopulation. For instance, reduced levels of the Treg suppressor marker CD39 have been reported in multiple sclerosis [9], systemic lupus erythematosus [10], type I diabetes mellitus [11], autoimmune hepatitis [12], rheumatoid arthritis [13] and asthma [14].

CD39 and HLA-DR are key components in Treg suppressive machinery. CD39, or ecto-nucleoside triphosphate diphosphohydrolase 1 (E-NTPDase1), is an enzyme of the ATPase family that hydrolyses ATP into AMP, which is subsequently hydrolysed by CD73 into adenosine [15,16]. CD39 plays a central role in modulating the ATP-driven pro-inflammatory microenvironment into an anti-inflammatory milieu controlled by the ability of adenosine to augment the function of regulatory T-cells [15] as well as to inhibit effector T-cell development [17]. Huge effort has recently been directed toward deciphering the TCR signature of T-cell clones that are involved in many autoimmune diseases, providing novel insight on the disease mechanisms, as TCRs are the main determinants of epitope specificity and phenotype predominance. Advances in sequencing technology allow in-depth analyses to be performed on the TCR repertoire of many subjects at the same time.

Antigenic epitopes are presented to TCRs in conjugation with major histocompatibility complex (MHC) molecules. Each unique TCR is generated by TCRα and TCRβ chain rearrangements, where the great amount of diversity found in the TRCβ chain particularly in the CDR3 region. CDR3 is generated by rearrangement of three genes: V, D and J, with a possibility of N nucleotide insertions in VD and DJ junction sites increasing diversity [18–21] Due to the vast number and diversity of the naïve T-cell repertoires, the small number of genes encoding for TCR have the potential to generate vast numbers of clonotypes, estimated at between 10^15^ and 10^20^. Although the process of recombination is thought to be random, there are some clonotypes that are common and these are known as public clonotypes [22].

## Results

Within the CD4+ T-cell pool, total Tregs were found to be 40% higher in peripheral blood of AA patients when compared to controls (P=0.001). Interestingly, a higher proportion 15% of CD45RO+ memory cells in the CD25+FOXP3+ Treg pool from patients (AA) was noted when compared to control (HC) samples (10%) (Figure 1A and Sup figure 1). The mean frequency of the suppressive Treg subset (CD39+ cells) was 43.5% of the CD25+FOXP3+ population in HC, however, this was significantly less in AA patients at 21.4% (P=0.001). There was also a reduction in expression of the HLA-DR suppressive marker on the surface of CD25+FOXP3+ cells with 38% of Treg HLA-DR positive in controls compared to 22% in AA patients (P≤0.0001). On the other hand, there was no change in the percentage of LAG-3+ cells in the Treg pool (Figure 1B and Sup figure 1). The reduction of CD39+ve PBMC in AA patients was confirmed by western blot. Fig 1C shows that expression of CD39 was greatly reduced in AA patients compared to controls, also when corrected by a semi-quantitative method (densitometry) using an internal control (GAPDH) to obtain relative expression (Figure 1C). To test whether the deficiency of circulating CD39+ Tregs associated with AA is also observed in CD39+ Treg cells resident in skin, we analysed the distribution of the suppressive Treg cell population in normal and AA skin. Suppressive Treg-specific immunofluorescence microscopic analyses were performed on 5 μm formalin-fixed, paraffin embedded sections obtained from healthy controls, as well as lesional and non-lesional areas of AA patients. Antibodies against CD3, FOXP3 (Treg marker) and CD39 (suppressive Treg marker) were used. Three populations were identified using this panel; CD3+FOXP3+CD39+ cells (suppressive Tregs), CD3+FOXP3+CD39-cells (non-suppressive Tregs that can be naïve or memory cells), and CD3+FOXP3-CD39-(T effector or Teff) cells. Consistent with the flow cytometry findings, we observed that suppressive Tregs in human skin preferentially localized to normal and non-lesional hair follicles (HFs) (Figure 2A). CD3+FOXP3+CD39+cells (suppressive Tregs) infiltrate the outer connective tissue layer of HFs. In contrast, diseased HFs lack these suppressive Tregs but show infiltration by CD_3_+FOXP_3_-CD_39_-conventional or effector T-cells (Tconvs/Teffs) with the presence of some generic Tregs lacking CD39 expression (CD3+FOXP3+ CD39-Tregs) (Fig 2A). Semi-quantitative analysis of HF images was performed to determine the frequency of Tregs and their suppressive subtype in normal, non-lesional and lesional skin. Interestingly, the proportion of total FOXP3+ Tregs was significantly reduced by about 80% in lesional compared to non-lesional skin (P≤0.01) and by about 90% when compared to normal skin (P≤0.001) (Figure 2B). A similar trend was observed with suppressive CD3+ Tregs where the proportion is significantly less in AA affected skin compared to non-lesional (P≤0.5) and normal skin (P≤0.001). The data is representative of the skin sections of lesional skin from four AA patients and non-lesional skin of three patients and one healthy control. Only anagen HFs were included in the analysis as AA is primarily a disease of the anagen HF, and to avoid the possibility that the variations observed in the sections were due to normal changes seen in the hair cycle stages.

**Figure 1.**
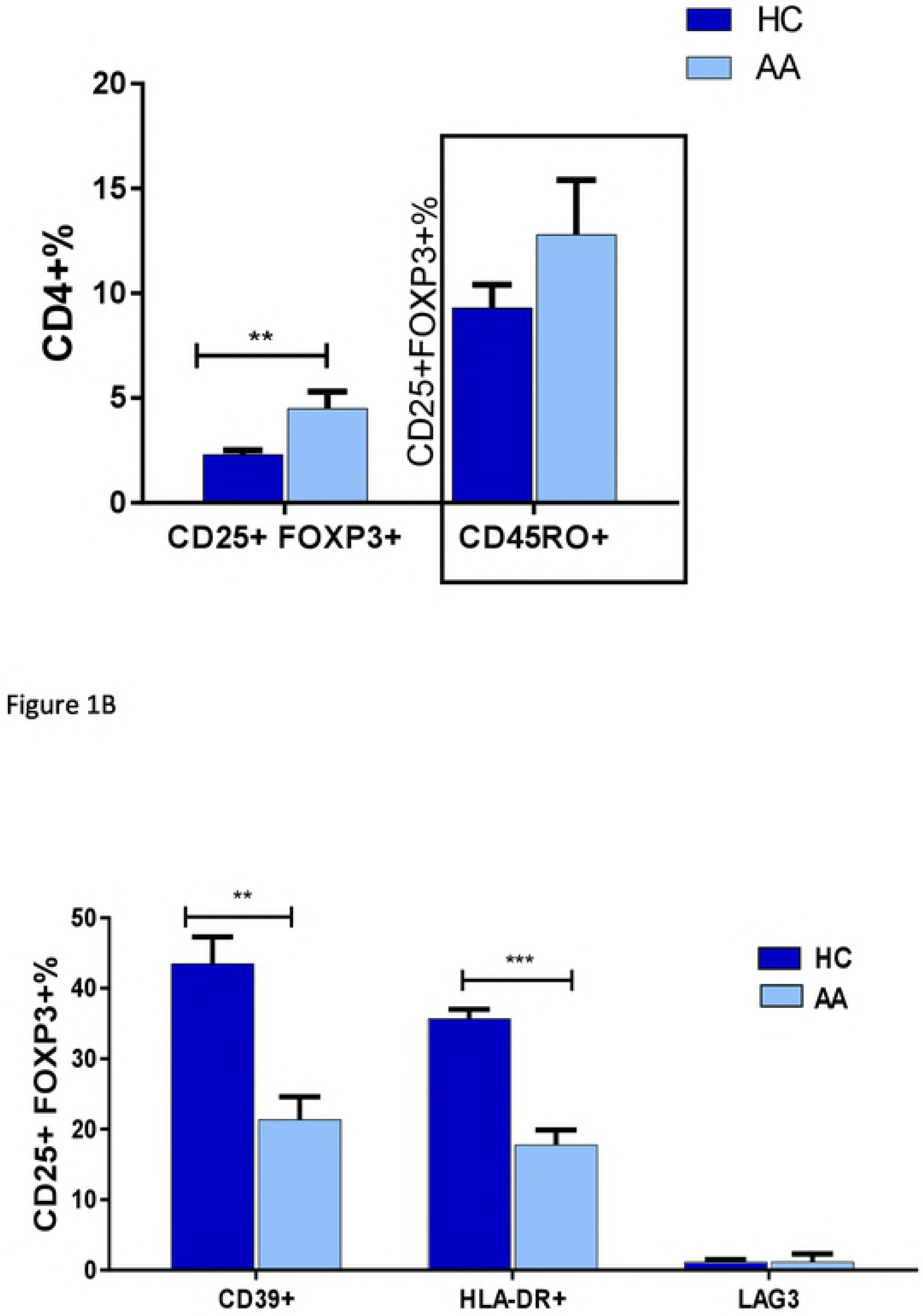
Flow cytometry identification strategies for Treg subsets (representative plots from one patient and one healthy control). **A:** Percentages of CD25+FOXP3+ Tregs and their memory subset (CD45RO+) in peripheral blood of AA patients and HC (see also supplementary data online). **B**: Suppressive subsets of CD25+FOXP3+ Treg cells. The percentage of suppressive subsets indicated by expression of CD39, HLA-DR or LAG3 surface makers was calculated out of the total Treg pool. Data analysed by a two-tailed independent t-test, and the corrected t-test was used whenever the homogeneity of variance was violated. A 95% confidence interval was used where P≤ 0.05 is considered significant (*) P≤o.o1 (**), P≤0.001 (***). Error bars depict SEM in each study group. AA patients n=20, healthy controls (HC) n=15. **C**: CD39 protein expression in AA and HC PBMCs. PBMCs isolated from blood of AA and HC donors were lysed and loaded in SDS gel, CD39 protein was detected using rabbit monoclonal antibodies. CD39 protein (60KDa) was detected in HC but not AA samples.

**Figure 2.**
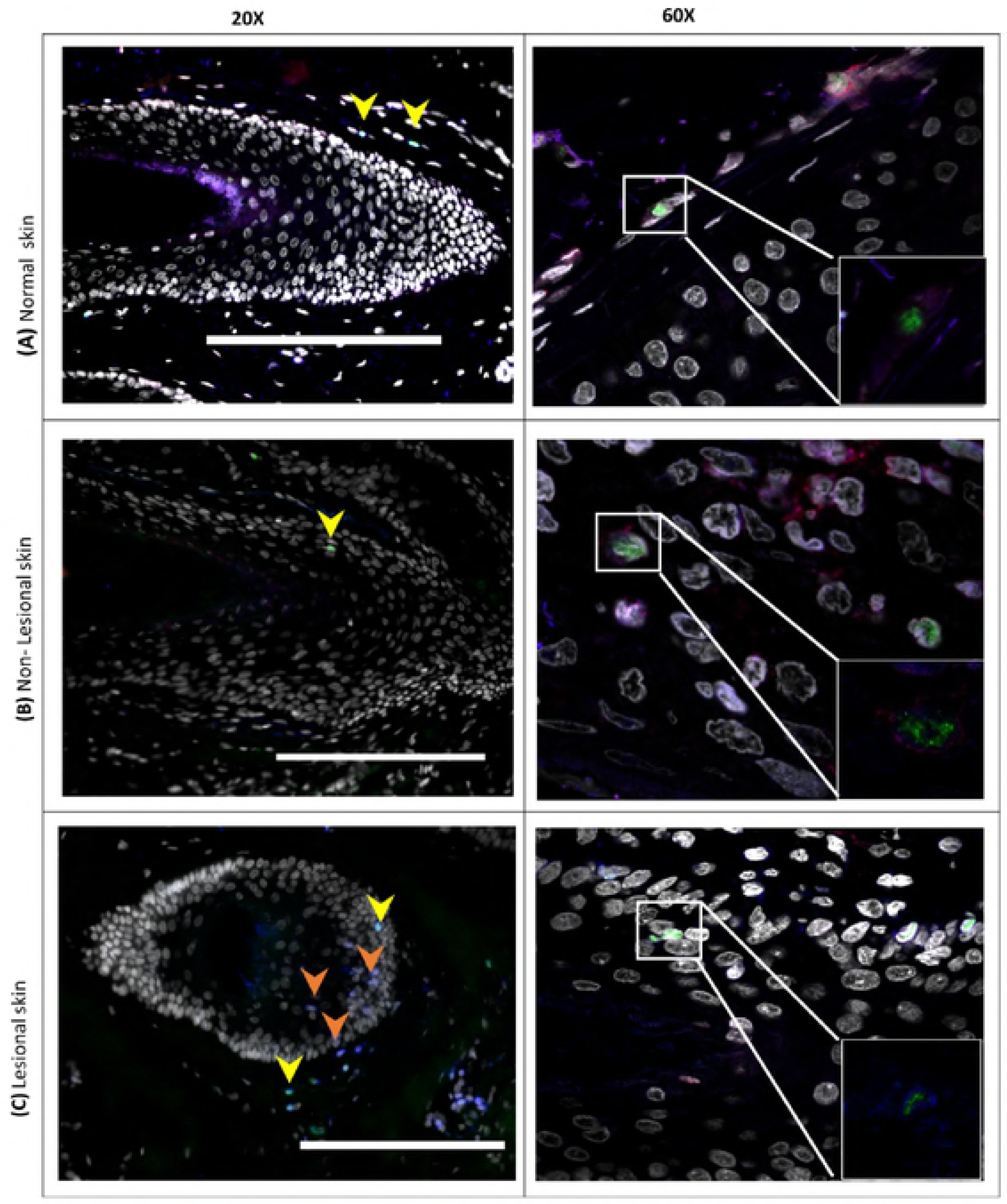
Suppressive Tregs localize to HFs in human skin. **A**. Confocal microscopy of normal human skin, lesional and non-lesional skin of AA patients. Sections were stained for CD3 (blue), FOXP3 (green), CD39 (red), and DAPI (gray). Images taken at magnifications X20 (left side) or X60 (right side). Small box (bottom right) is the zoom on 60X images without nuclei to clarify the triple staining. Scale bars: 100 μm. Arrows denote CD3+FOXP3+CD39+ cells or CD3+FOXP3+. The data is a representation of the skin section of one AA patient and one healthy control. We looked at three patients, which showed the same trend in suppressive Treg distribution (data not shown). **B**. Suppressive subsets of CD25+FOXP3+ Treg cells. The percentage of suppressive subsets indicated by its expression of CD39 and FOXP3 makers was caluclated out of the total CD3+ T-cell pool. Data are derived from the lesional skin of four AA patients, non-lesional skin of three patients and one healthy control. Data from of four fields in each study group were analysed by ANOVA with multiple comparison. A 95% confidence interval was used with *P≤* 0.05 (after a Bonferroni correction) is considered significant (*), P≤0.001 (***). Error bars depict SEM.

To further characterise Tregs in AA, Tregs (CD4+CD25+FOXP3+) FACs sorted from PBMCs of patients and controls were investigated for their TCRβ sequence characteristics as described in materials and methods section. The TCR repertoire of total lymphocytes from AA patients was slightly less diverse (chao1=3.2) compared to controls (chao1=3.4), which was likely due to the presence of dominant clonotypes. However, the Treg repertoire showed a higher diversity in patients (chao1=2.4) when compared to controls (chao1=2.3) reflecting the predominance of unique clonotypes of Treg cells in HC (Figure 3). The public TCR sequence shared by many individuals suggests the exposure of those individuals to the same antigen. The number of unique CDR3 clonotypes in the TCRβ repertoire was 1.8×10^5^ in PBMCs of AA patients, 1.51×10^5^ in PBMCs of HC, 1.07×10^5^ in Tregs in AA patients and 0.8×10^5^ in Tregs of HC. The most frequent public clonotype in each group is presented in figure 4. This figure shows that the most frequent public clonotypes in AA Tregs (FACS AA) were also present in controls (FACS HC) at relatively similar frequencies, except for CASTKTKRQGPISRPFPTGELFF (V12-3 D1 J2-2) CANSTRGS_PGNTIYF (V10-1 D2 J1-3) and CASSPTGPTEAFF (V7-2 D1 J1-1) which were more abundant in Tregs isolated from patients (with frequency of 0.02, 0.01 and 0.08 respectively) compared to controls (0.008, 0.002 and 0.003). However, there were two public clonotypes CATSRDEGGLDEKLFF (V15 D1 J1-4) and CASRDGTGPSNYGYTF (V2 D1 J1-2) that were exclusively represented in controls Tregs but not in patients. In contrast, two relatively frequent public clonotypes CASSYQGSTEAFF (V6-3 D1 J1-1) and CASSQDKGITNEKLFF (V4-2 D1 J1-4) were exclusively represented in AA PBMC but not in controls. These were not Tregs because not present among AA Treg clonotypes. When comparing V segment usage by Tregs in patients and controls we found that TRBV12-4, 20-1, 24-1 were more predominant in cases compared to controls, while other TRBV segments such as TRBV12-3, 2, 30 and 15 were highly abundant in controls but not in cases. Applying post-hoc statistical analysis, the frequency of TRBV2 segments was significantly higher in controls compared to patients (P=0.01), on the other hand, TRBV 6-3 showed a significant increase in its frequency in Tregs of patients (P=0.005) (Figure 5). Similarly, when looking at J segment usage, TRBJ1-2 and to less extent TRBJ2-5 and TRBJ2-7 were slightly more abundant in cases. Controls have more abundant TRBJ2-2, 1-5, and 1-1 compared to cases. The frequency of TRBV 1-1 usage is significantly higher in controls (P=0.04), and TRBV2-5 is higher in patients (P=0.05) (Figure 6). The CDR3 nucleotide (nt) length ranged from 21 to 69nt, with most frequently observed sizes of 39, 42, 45 and 48 nucleotides in TCRβ chains of total lymphocytes as well as Tregs in both patients and controls (Figure 6A and B). Similarly, the NDN nucleotide size distribution was identical in Tregs of patients and controls with 13nt the most frequent size. In total lymphocytes, NDN length showed a similar distribution ranging from 0 to 40nt; however, the most frequent length in controls was 13nt while it was 9nt in patients. Looking at the insert between VD and VJ regions, a similar length distribution can be observed in lymphocytes, where no insert (0nt) was the most commonly observed length in both groups followed by 4nt length in patients and 7nt length in controls (Figure 6C). In contrast, a marked difference can be seen in VD/VJ indel (insertion/deletion) length in Tregs; while no insert (0 nt) is the most frequent distribution in patients, the inserts predominantly found in VD/VJ region in controls with about 13nt indel the most commonly observed length, which may indicate the presence of a base deletion at the V-D/V-J junction in AA patients. No significant difference was observed between NDN nucleotide size between patients and controls when considering total PBMC (Sup figure 2).

**Figure 3.**
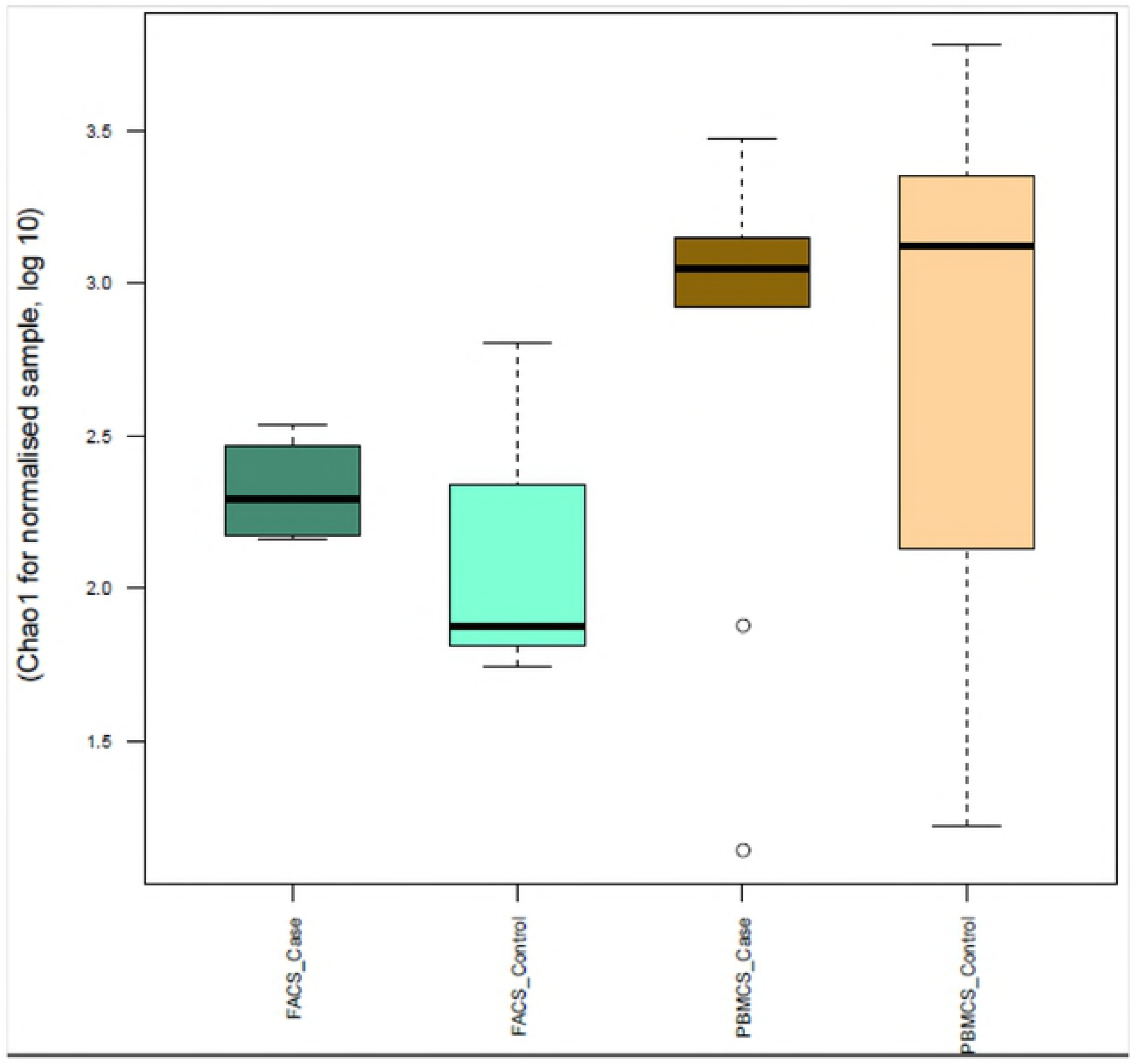
TCR Repertoire diversity estimation. Boxplot of chao1 estimator for normalised samples were used to assess the diversity of TCR repertoire in cases and control. PBMC: total blood cells. FACs: Sorting for Treg cells by FACs analysis.

**Figure 4.**
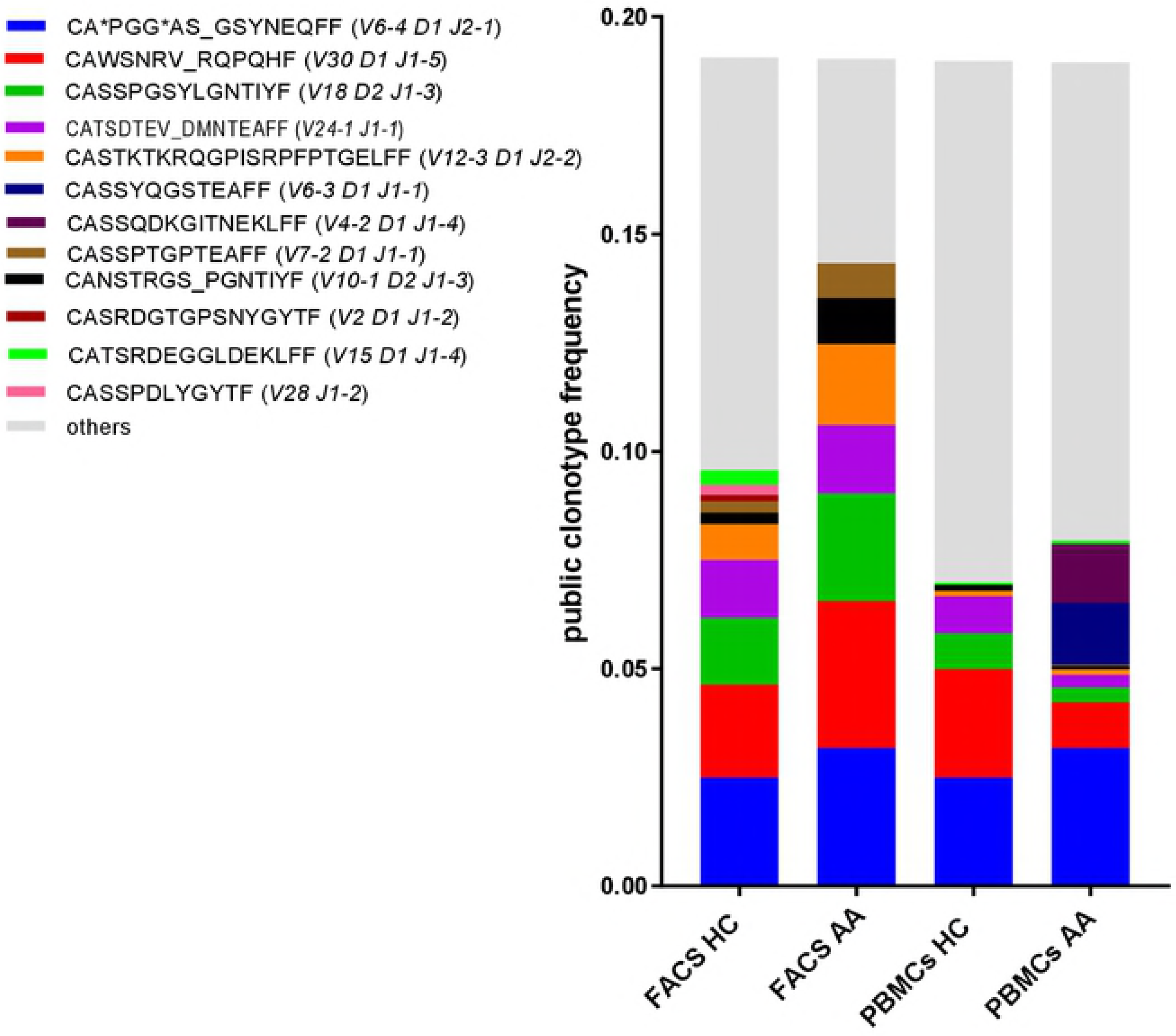
Distribution of public clonotypes in Tregs and total T-cells of patients and HCs. The most frequent sequences are allocated individual colours defined in the key, while the others are combined and shown in grey. PBMC: total blood cells. FACs: Sorting for Treg cells by FACs analysis.

**Figure 5.**
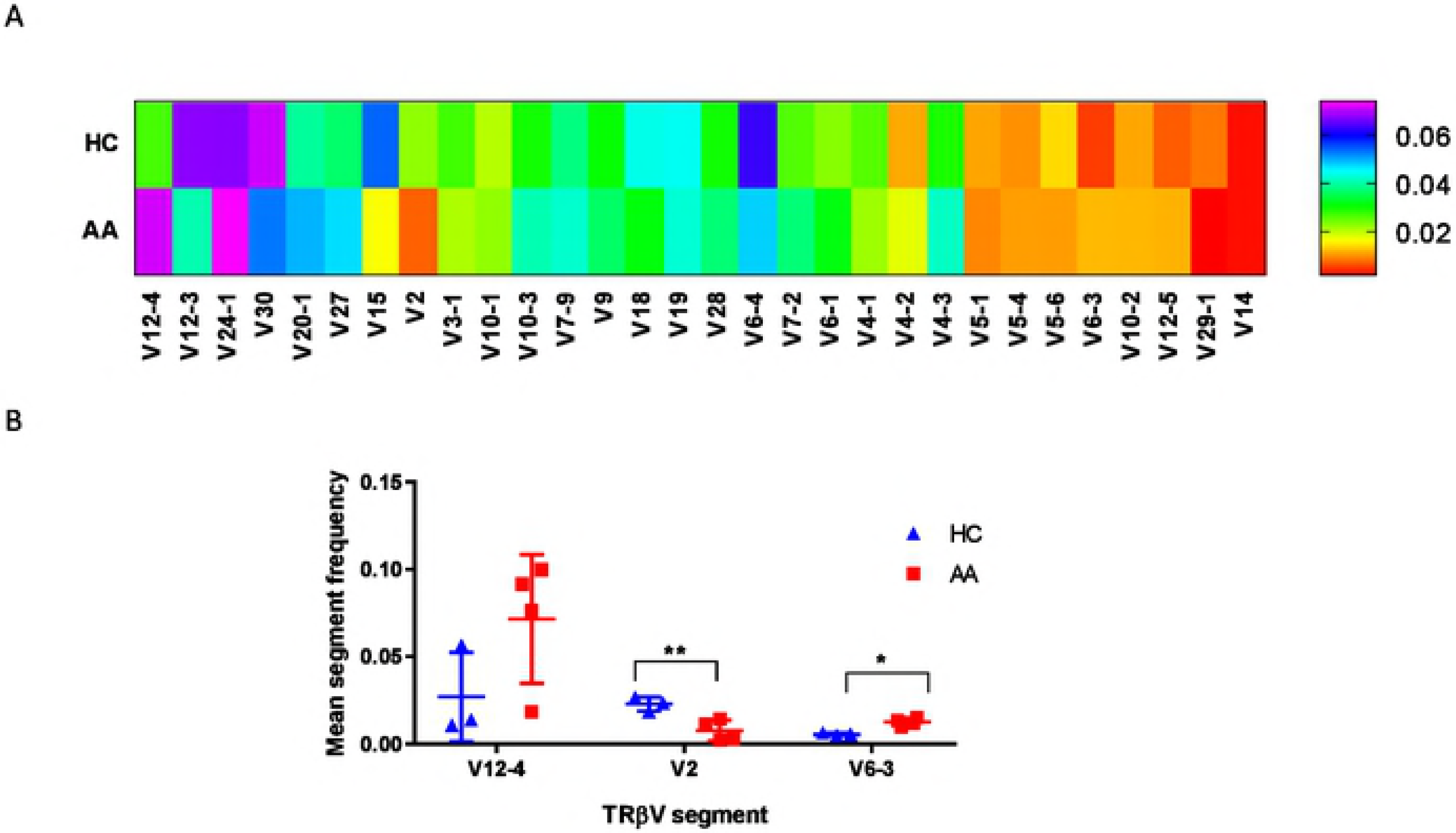
Gene segment usage in TCR of Tregs. **A**. frequency of each V segment usage in healthy control (HC) and AA cases, the data represented in the heat map is the mean of three HCs and four AA patients **B**. Statistical analysis of the observed changes in TCRβV segment usage between HC and AA where the difference was calculated by ANOVA and * P<0.05.

**Figure 6.**
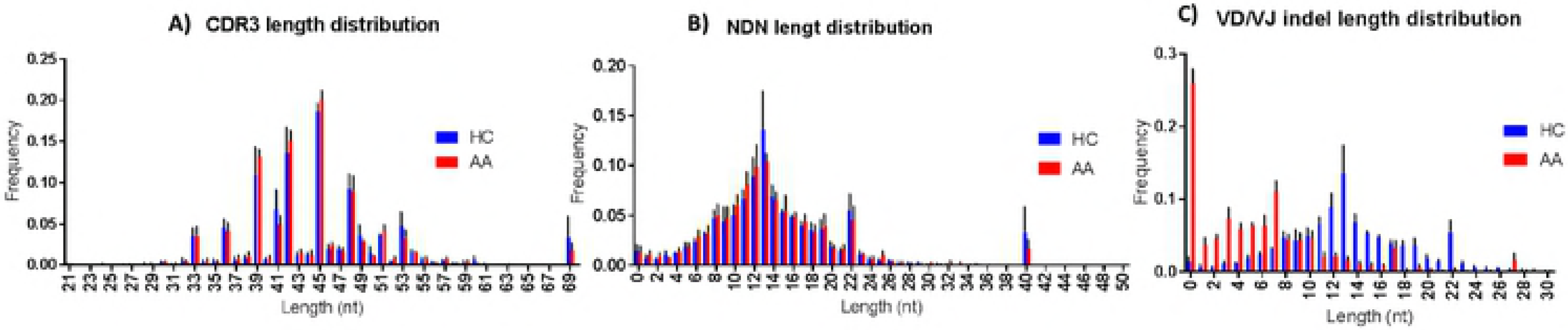
CDR3 bulk characteristics in Treg of patients and healthy controls. **A)**. nucleotide length of CDR3 segment **B)**. NDN nucleotides size **C)**. V-D and D-J insert size. NDN and CDR3 length in cases and controls showed same distribution for Treg and total lymphocytes.

## Discussion

In this study we report, for the first time, that the distribution of Tregs in lesional AA tissue compared to healthy and non lesional tissue and show there are marked abnormalities in Treg subsets in patients with AA. The reduction of circulating CD39+ Tregs of AA patients was in keeping with the reduction of these cells in the skin of AA patients, and suggests that the flow of CD39+ Tregs into the skin, making the HF vulnerable to autoimmune attack. In this respect, the significant reduction of CD39+ Tregs in the HF and its surrounding area of uninvolved skin could be a landmark of predisposed non-lesional skin to become alopecic. This is in keeping with Castela and colleagues’ study showing that recruitment of Tregs in lesional skin promotes hair regrowth in AA patients [23]. An intriguing observation regarding the potential importance of CD39 deficiency in the pathogenesis of AA comes from a study of CD39 knockout mice [24] in which it was documented that 15% of CD39 null mice developed spontaneous autoimmune alopecia characterised by lymphoid infiltration of damaged hair follicles and a 6-fold increase in interferon-gamma transcript levels in areas of skin with active disease, in comparison with uninvolved or normal control skin. Our novel finding in the present study, of decreased CD39 expression on T-cells in patients with active hair loss, is an interesting addition to the understanding of the immunology of AA and raises the possibility that the disease may be amenable to therapeutic approaches involving reconstitution of CD39 +ve Tregs. CD39+ deficiency has also been recently shown in psoriasis with a significant reduction of CD39+ FOXP3+ cells observed in patients with erythrodermic and pustular psoriasis [25]. Our flow-cytometry data also demonstrated significant reduction of HLA-DR+ Tregs and HLA-DR expression on Tregs mediates cell-to-cell suppression of Teffs. Interestingly, Rissiek et al (2015) showed that HLA-DR expressing Tregs are almost always CD39+, which could explain the reduction of Tregs HLA-DR+ cells in AA patients as observed in the current study [26]. The suppressive cytokine secretory capacity of Tregs was investigated and showed no significant changes in the frequency of IL-10+ nor TGF-β+ Tregs between patients and HC (Sup Figure 2). These findings are supported by LAG-3 expression, which showed no difference between the groups (Figure 1B). The activation gene (LAG3) has been reported as a marker of IL-10 producing Treg [27]. Taking these findings together suggests that impaired Treg function in AA is mainly due to a defect in cell-to-cell contact and CD39-mediated suppressive machinery and not in cytokine secretion.

In this study, CASSYQGSTEAFF (V6-3 D1 J1-1) and CASSQDKGITNEKLFF (V4-2 D1 J1-4) were only found in the TCR repertoire of total lymphocytes of patients, suggesting that cells using these two public clonotypes are not Tregs (not detected in AA Tregs) and maybe involved in AA pathogenesis as they were not detected in controls. Database searches showed that CASSYQGSTEAFF has 86% match to CDR3 of peripheral T-lymphocytes of Sjögren’s syndrome [28] whereas CASSQDKGITNEKLFF sequence has about 81% similarity was found with TCRβ chain from autoreactive CD8+ T-cells isolated from diabetic mice [29]. The TCR repertoire of Tregs showed that two TCRβ clones CATSRDEGGLDEKLFF (V15 D1 J1-4) and CASRDGTGPSNYGYTF (V2 D1 J1-2) in healthy controls were totally absent in patients, suggesting that these Treg clonotypes are protective. Such cells could potentially be expanded *in vitro* and used for treatment. In fact, a single clone of Treg has been recently expanded *in vitro* and was successfully used to reduce inflammation and neovascularisation in diabetic retinopathy in mice model [30], suggesting that deciphering the clonotypic map of T-cells in AA would bring the field a step closer toward understanding the disease pathogenesis.

## Materials & Methods

### 1. Peripheral blood lymphocytes

#### 1.1 Blood samples

The study was approved by the Institutional Review Boards/ Ethics Committees at the University of Sheffield (LREC reference number 002651) and Sheffield Teaching Hospitals (STH permission reference number STH18941). Twenty patients with active hair loss and an established diagnosis of AA were recruited and consented at the Department of Dermatology, Royal Hallamshire Hospital, Sheffield, UK. Patients diagnosed with other autoimmune diseases or receiving immunosuppressive drugs were excluded from the study. The cases recruited included 9 with patchy AA, 5 with alopecia totalis, and 6 with alopecia universalis. Patients and healthy controls were age-matched and were all female of Caucasian ethnicity.

#### 1.2. Peripheral blood mononuclear cell separation and FACS analysis

Peripheral blood mononuclear cells (PBMC) were isolated from heparinized venous blood by density gradient purification using Lymphoprep as described by the manufacturer (07801, Stem Cell) and stained with a panel of antibodies to Treg markers. Tregs were identified by co-expression of CD25 and FOXP_3_ in CD4+ cells, and Treg subtypes including memory CD4+CD25+FOXP3+ and effector/suppressive CD4+CD25+FOXP3+CD39+/ CD4+CD25+FOXP3+HLA-DR+/CD4+CD25+FOXP3+LAG-3+ were defined by the surface markers CD45RO, CD39, HLA-DR and LAG-3 (Sup Tables 1). Briefly, 10^6^ PBMC were incubated for 30 minutes at room temperature with blue fixable live/dead dye (Thermo-Fisher), washed once with phosphate buffered saline (PBS) and stained with antibodies (BD, UK) targeting surface markers in each panel. The cells were then fixed and permeabilised by fix/perm buffer (Transcription buffer set, 562725, BD) for 40-50mins at 4°C, followed by intracellular staining by incubation for 40-50mins at 4°C with antibodies specific for markers above. The cells were finally fixed in 2%PFA and the staining visualised by LSR II (Becton Dickinson, Heidelberg, Germany). Further gating was performed by Flow Jo software to determine the frequency of T-cell subpopulations. Gating of the positive population for each marker was performed based on Fluorescence minus one control (FMO). Unstained, single cell controls and compensation controls were also used to optimise the experimental parameters.

#### 1.3 Immunoblot analysis

PBMC cell lysates were prepared by lysis in RIPA buffer (150 mM sodium chloride, 50mM Tris-HCl, pH 7.4, 2 mM ethylenediaminetetraacetic acid, 1% Triton X-100, 0.5% sodium deoxycholate, 0.1% sodium dodecylsulfate) containing protease inhibitor cocktail (P8340-5ML, Sigma). Cell debris was removed by centrifugation. Protein concentration was determined by BCA assay (Pierce™ BCA protein assay kit, 23225, Thermofisher). Lysates were then boiled for 5 min in 1 × SDS sample buffer (B31010, Lifetechnologies) and proteins separated by SDS-PAGE (pH8.8, 10% [37:1] acrylamide, 0.375M Tris-Cl and 0.1% SDS). After electrophoresis, the gel was transferred to a PVDF membrane using the iBlot system (Invitrogen). Primary monoclonal antibodies used were anti-CD39 IgG (rabbit, ab108248, Abacam, 1:1000) and anti-GAPDH IgG (rabbit, ab128915, Abcam, 1: 10,000). The secondary antibody was 1:10,000 goat anti-rabbit IgG conjugated with peroxidase (4050-05 Southern Biotech). The membrane was developed using EZ-ECL chemiluminescent reagent (Biological Industries) and visualised using the ChemiDoc XRS+ System (Bio-Rad).

#### 1.4 Statistical analysis

The percentage of each T-lymphocyte subpopulation was compared between patients and controls using a two-tailed independent t-test and ANOVA. The analysis was done using SPSS version 22 with *P* < 0.05 as the significance level (see supplementary data).

### 2. Human tissue samples

#### 2.1 Preparation of tissue sections

Lesional and non lesional scalp skin biopsies were obtained from AA patients after patient consent and ethical approvals from the University ‘La Sapienza’ of Rome, Italy (n.2973, 28-11-13) and of the University of Münster, Germany (2014-041-b-N, and 2015-602-f-S). The samples were fixed in 4% formalin and embedded in paraffin. Sections of 4.5μm were prepared. Normal skin was obtained from healthy control.

#### 2.2 Immunofluorescence microscopy

An immunofluorescent detection technique was used to co-localise CD3, CD39 and FOXP3 as markers of suppressive Tregs in hair follicle (HF) in formalin-fixed, paraffin-embedded (FFPE) skin tissue from AA and healthy control. Antigens were retrieved from deparaffinised sections by heating in a microwave set to full power for 10 mins. Non-specific binding sites were blocked using serum-free blocking buffer. The target antigens were initially labelled with primary antibodies to surface antigens (CD3 or CD39) overnight at 4°C followed by labelling with the primary antibody against the intracellular marker for 2hrs at room temperature in 0.2% Triton X in PBS to facilitate membrane permeabilisation. The sections were then incubated with fluorochrome-conjugated secondary antibody for 1hr at room temperature, and the signal detected by immunofluorescence-scanning microscopy (Leica AF6000). Antibodies are listed in the supplementary data (Sup Table 3). The number of CD3+, FOXP3+ and CD39+ cells was counted per field, and the ratio of FOXP3+ cells to total CD3+ T-cells in each field was calculated using FUJI cell counter software. The ratio of CD39+ cells in the FOXP3+ cell pool was calculated by the same method.

### 3. Next Generation Sequencing

The detailed protocol used for Next Generation Sequencing (NGS) is in supplementary materials. Briefly, DNA was extracted from 10^6^ PBMCs from 14 AA patients and 7 healthy controls. PBMCs were isolated from heparinized venous blood by density gradient purification as described in section 2.2.2, and 10^4^ CD4+CD25+FOXP3+ Tregs sorted by FACS technique. For multiplex-PCR amplification of the *TRCB* CDR3 region, TCRβ CDR3 was defined according to the International Immunogenetic Collaboration whereby TCRβ CDR3 begins with the second conserved cysteine encoded by the 3’ position of the Vβ gene segment and ends with the conserved phenylalanine encoded by the 5’ position of the Jβ gene segment. A multiplex-PCR system was used to amplify the rearranged regions of TCRβ CDR3 from genomic DNA using 39 forward primers specific to TCR Vβ segments and 13 reverse primers each specific to a TCR Jβ segment to generate a template library using Genome Analyzer. DNA fragments (multiplex PCR product of PBMCs samples) around 500bps were fragmented into about 200-300bp size after 20mins incubation with fragmentase enzyme at 37°C. Bead purification of the PCR products was performed as described by the supplier AgenCourt AMPure XP (A63881, Beckman Coulter) to remove excess enzymes, primers, salts and nucleotide. Sequencing libraries were generated using the standard protocol in the NEB Ultra II DNA library prep kit for Illumina (E7645S, NEB).

Multiplex PCR was performed using 39 V primers and 13 J primers to amplify the rearranged CDR3 region and the PCR products analysed by NGS technique using the Illumina platform. Base calling accuracy was measured by the Phred quality score (Q score), which calculates the probability of calling the incorrect base, and 89% of the data was above Q30 indicating the probability of calling the right base was 99.9% (Sup Figure 2). To determine the impact of clonal expansion on the diversity of the TCR repertoire in total lymphocytes and Tregs of AA patients, biodiversity was evaluated in each group using a chao1 normalised sample estimate. To investigate clonotype overlap, VDJtools produced pool strict data (pooled clonotype abundance table with the columns as described at http://vdjtools-doc.readthedocs.io/en/latest/operate.html), where the number of reads associated with public clonotypes was computed in each group as well as their frequency. Public clonotypes were defined as those having identical CDR3 nt sequence, VDJ arrangement and start & end position for each segment.

## Conflict of Interest

The authors state no conflict of interest.

## Acknowledgment

We thank Prof Alison Condliffe for her comments and constructive criticism, and Mrs Kathy Dobson for recruiting alopecia areata patients who kindly volunteered to participate in this study. We also thank Dr. Paul Heath in the Sheffield Institute for Translational Neuroscience (SITraN) for library preparation for NGS. Financial support was provided under the Contract Research Organization agreement between AstraZeneca and University of Sheffield. FNH was awarded a studentship (no. 313821 MED) from the Libyan Ministry of Education.

## Supplementary material

Supplementary material is attached to this manuscript

